# Fibroblasts and Alectinib switch the evolutionary games played by non-small cell lung cancer

**DOI:** 10.1101/179259

**Authors:** Artem Kaznatcheev, Jeffrey Peacock, David Basanta, Andriy Marusyk, Jacob G. Scott

**Affiliations:** Department of Computer Science, University of Oxford; Department of Translational Hematology & Oncology Research, Cleveland Clinic; Department of Radiation Oncology, Moffitt Cancer Center; Department of Integrated Mathematical Oncology, Moffitt Cancer Center; Department of Cancer Imaging and Metabolism, Moffitt Cancer Center; Department of Radiation Oncology, Cleveland Clinic

## Abstract

Heterogeneity in strategies for survival and proliferation among the cells which constitute a tumour is a driving force behind the evolution of resistance to cancer therapy. The rules mapping the tumour’s strategy distribution to the fitness of individual strategies can be represented as an evolutionary game. We develop a game assay to measure effective evolutionary games in co-cultures of non-small cell lung cancer cells which are sensitive and resistant to the anaplastic lymphoma kinase inhibitor Alectinib. The games are not only quantitatively different between different environments, but targeted therapy and cancer associated fibroblasts qualitatively switch the type of game being played by the in-vitro population from Leader to Deadlock. This observation provides empirical confirmation of a central theoretical postulate of evolutionary game theory in oncology: we can treat not only the player, but also the game. Although we concentrate on measuring games played by cancer cells, the measurement methodology we develop can be used to advance the study of games in other microscopic systems by providing a quantitative description of non-cell-autonomous effects.

---

Tumours are heterogeneous, evolving ecosystems [1, 2], comprised of sub-populations of neoplastic cells that follow distinct strategies for survival and propagation [3]. The success of a strategy employed by any single neoplastic sub-population is dependent on the distribution of other strategies, and on various components of the tumour microenvironment, like cancer associated fibroblasts (CAFs) [4]. The EML4-ALK fusion, found in approximately 5% of non-small cell lung cancer (NSCLC) patients, leads to constitutive activation of oncogenic tyrosine kinase activity of ALK, thereby “driving” the disease. Inhibitors of tyrosine kinase activity of ALK (ALK TKI) have proven to be highly clinically efficacious, inducing tumour regression and prolonging patient survival [5, 6]. Unfortunately, virtually all of the tumours that respond to ALK TKIs eventually relapse [7] – an outcome typical of inhibitors of other oncogenic tyrosine kinases [8]. Resistance to ALK TKI, like most targeted therapies, remains a major unresolved clinical challenge. Despite significant advances in deciphering the resultant molecular mechanisms of resistance [9], the evolutionary dynamics of ALK TKI resistance remains poorly understood. The inability of TKI therapies to completely eliminate tumour cells has been shown to be at least partially attributable to protection by aspects of the tumour microenvironment [10]. CAFs are one of the main non-malignant components of tumour microenvironment and the interplay between them and tumour cells is a major contributor to microenvironmental resistance, including cytokine mediated protection against ALK inhibitors [11].

To study the eco-evolutionary dynamics of these various factors, we interrogated the competition between treatment naive cells of ALK mutant NSCLC cell line H3122 – a “workhorse” for studies of ALK+ lung cancer – and a derivative cell line in which we developed resistance to Alectinib – a highly effective clinical ALK TKI [12] – by selection in progressively increasing concentrations of the drug [13]. We aimed to come to a quantitative understanding of how these dynamics were affected by clinically relevant concentrations of Alectinib (0.5µM; see [14]) in the presence or absence of CAFs isolated from a lung cancer. To achieve this, we developed an assay for quantifying effective games [15, 16] that is of independent interest to the general study of microscopic systems.

## Results

### Monotypic vs mixed cultures

To establish baseline characteristics, we performed assays in monotypic cultures of parental (Alectinib-sensitive) and resistant cell lines with and without Alectinib and CAFs. To gather temporally-resolved data for inferring growth rates, we used time lapse microscopy to follow the expansion of therapy resistant and parental cells, differentially labeled with stable expression of selectively neutral GFP and mCherry fluorescent proteins, respectively. From the time series data, we inferred the growth rate with confidence intervals for each one of 6 experimental replicates in four different experimental conditions (total of 24 data points, each with confidence intervals), as seen in Figure 1. As expected, alectinib inhibited growth rates of parental cells (DMSO vs Alectinib: *p* < .005; DMSO + CAF vs Alectinib + CAF: *p* < .005), whereas the growth rate of the resistant cells was not affected. And, as previously reported [11], CAFs partially rescued growth inhibition of parental cells by Alectinib (Alectinib vs Alectinib + CAF: *p* < .005; Alectinib + CAF vs DMSO: *p* < .005), without impacting growth rates of resistant cells.

**Figure 1:**
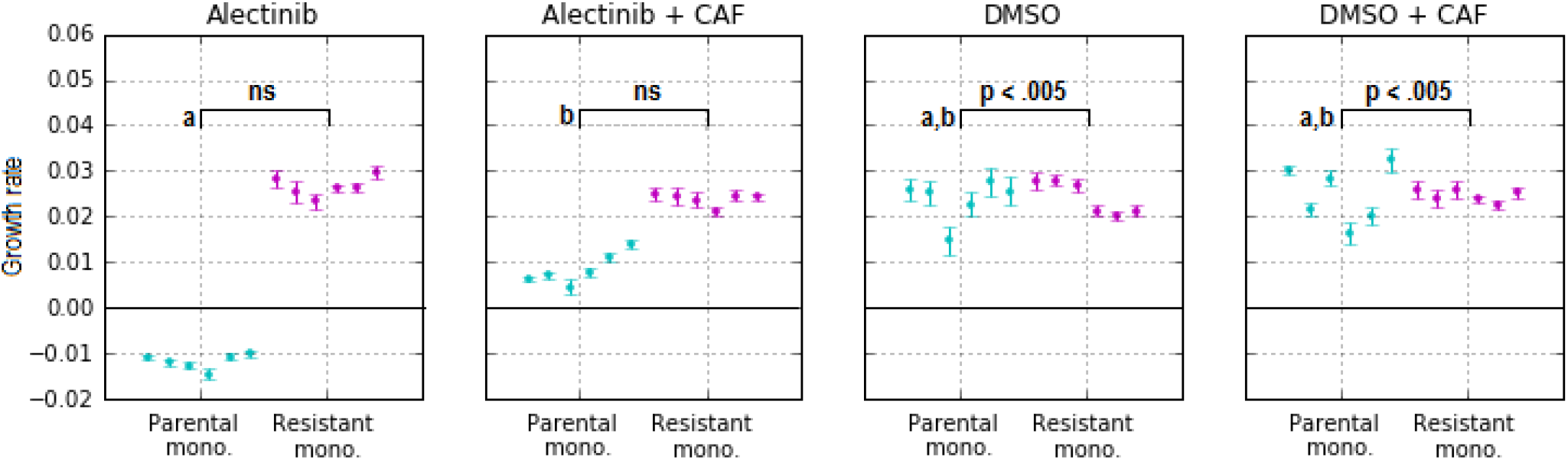
Monotypic culture exponential growth rates for parental (cyan) and resistant (magenta) cells in indicated experimental conditions. Confidence intervals on each experimental replicate is from confidence on the estimate of growth rate for that single replicate according to the Theil-Sen estimator. Comparisons between experimental conditions (of 6 replicates each) are made using Wilcoxon rank-sum. In addition to conditions linked by lines with reported p-values, conditions labeled by ‘a’ and ‘b’ are pairwise distinguishable with *p* < .005.

But we did not limit ourselves to monotypic assays. Our experience observing non-cell-autonomous biological interactions [17] and modeling eco-evolutionary interactions [18–20] in cancer led us to suspect that the heterotypic growth rates would differ from monotypic culture. Cell-autonomous fitness effects are ones where the benefits/costs to growth rate are inherent to the cell: the presence of other cells are an irrelevant feature of the micro-environment and the growth rates from monotypic cultures provide all the necessary information. Non-cell-autonomous effects [17] allow fitness to depend on a cell’s micro-environmental context, including the frequency of other cell types: growth rates need to be measured in competitive fitness assays over a range of seeding frequencies. Other microscopic experimental systems in which frequency dependent fitness effects have been considered include, but are not limited to: *Escherichia coli* [21, 22], yeast [23, 24], bacterial symbionts of hydra [25], breast cancer [17] and pancreatic cancer [26]. Hence, we continued our experiments over a range of initial proportions of resistant and parental cells in mixed cultures for each of the four experimental conditions.

Figure 2 shows the resulting growth rates of each cell type in the co-culture experiments for all experimental (color, shape) and initial conditions (opacity is parental cell proportion). In the heterotypic culture – unlike monotypic – CAFs slightly improved the growth rates of the parental cells, even in DMSO. More strikingly, even in the absence of drug, resistant cells tend to have a higher growth rate than parental cells in the same environment (i.e. proportion of parental cells in the co-culture). This is evident from most DMSO points being above the dotted diagonal line (*y* = *x*) corresponding to equal growth rate of the two types (this is quantified in Figure 4b and is further discussed in section ‘Leader and Deadlock games in NSCLC).

### Frequency dependence in fitness functions

Although not common in cancer biology, competitive fitness assays are a gold standard for studying bacteria. But they are typically conducted with a single initial ratio of the two competing cell types. However, in Figure 2, if we view the initial proportion of parental to resistant cells as a variable parameter represented by opacity then we can see a hint of frequency dependence in both parental and resistant growth rates. This is shown more clearly as a plot of fitness versus proportion of parental cells in Figure 3. In all four conditions, we see that the growth rate of the resistant and parental cell lines depends on the initial proportion of parental cells. To capture the principle first-order part of this dependence, we consider a line of best fit between initial proportion of parental cells and the growth rates. See equations 1-8 in Supplementary Appendix C (or the matrix entries in Figure 4b) for these lines of best fit. Interpretable versions of these lines of best fit (see Supplementary Appendix D) can be expressed as a regularized fitness function 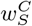 where *S* ∈ {*P, R*} indexes the parental or resistant strategy and *C* ∈ {DMSO, DMSO + CAF, Alectinib, Alectinib + CAF} indexes the experimental condition. For a description of regularization see Supplementary Appendix D. Finally, for a discussion of higher-order fitness functions, see Supplemental Appendix F.

In three of the conditions, resistant cell growth rates increase with increased seeding proportion of parental cells, while parental growth rates remain relatively constant (in the case of no CAFs) or slightly increase (for Alectinib + CAFs). In DMSO, this suggests that parental cells’ fitness is independent of resistant cells: 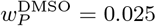. Parental fitness in DMSO could be well characterized as cell-autonomous. However, resistant cells in monotypic culture have approximately the same fitness as parental cells (Figure 2a), but they benefit from the parental cells in co-culture: 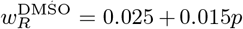 (where *p* is the proportion of parental cells). Their fitness has a non-cell autonomous component. The positive coefficient in front of *p* suggests commensalism between resistant and parental cells, i.e. resistant cells benefit from the interaction with the parental cells, without exerting positive or negative impact on them.

**Figure 2:**
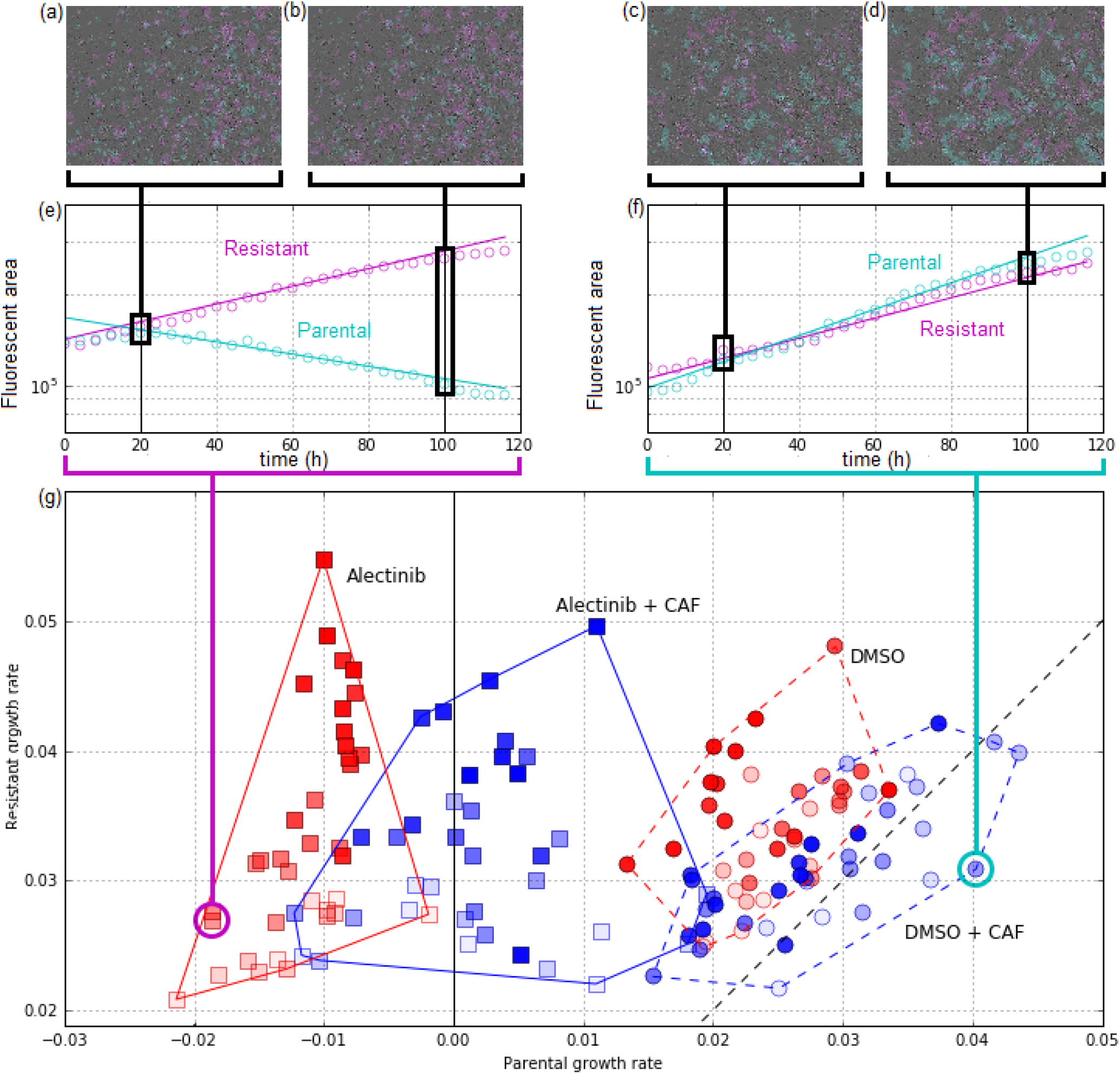
Coculture growth rates across four experimental conditions. (**a-f**) serve as a sketch of the analysis procedure to produce the main subfigure (**g**); for more detailed discussion, see Supplemental Appendix B.4 (**a,b,c,d**): In each experimental replicate at each time step, we quantify population size by fluorescent area of each cell type (shown: two different time points per well, from two different wells). Together, 30 time-lapse microscopy images (one every 4 hours) from each replicate create (**e,f**): time-series of parental and resistant population size (shown: two example wells). With x-axis is time, y-axis is log of population size. Exponential growth rates (and confidence intervals; omitted) were estimated for each well using the Theil-Sen estimator. These exponential models are shown as solid lines and their slopes serve as the coordinates in (**g**). See Figure 3 for growth rate confidence intervals and Supplemental Section B.2 for detailed discussion of growth-rate measurement. (**g**): Each point is a separate replicate of a competitive fitness assay with initial proportion of parental cells represented by opacity and experimental condition represented by shape (DMSO: circle; Alectinib: square) and colour (no CAF: red; + CAF: blue). Each replicate’s x-position corresponds to the measured parental growth rate and y-position for resistant growth rate; the dotted black line corresponds to the line of equal fitness between the two at *x* = *y*.

**Figure 3:**
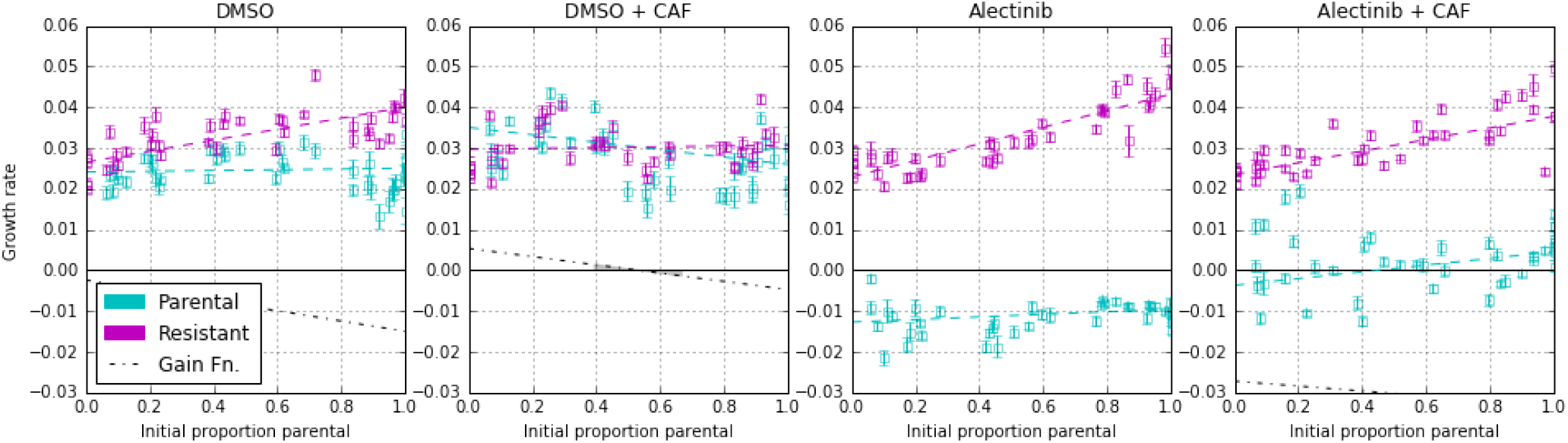
Fitness functions for competition of parental vs. resistant NSCLC. For each plot: growth rate with confidence intervals versus initial proportion of parental cells. This is the same data, measured in the same way, as Figure 2. Cyan data points are growth rates of parental cells, and magenta for resistant cells. Dotted lines represent the linear fitness function of the least-squares best fit; fit error is visualised in Figure 4b. The black dotted line is the gain function for parental (see Figure 4a), it is well below the *y* = 0 line in the Alectinib conditions (indicating the strong advantage of resistance) and thus cut out of the figure. See Supplemental Appendix C for more discussion and equations for lines of best fit, and Supplemental Appendix F for alternative fits with non-linear fitness functions.

The DMSO + CAF case differs from the other three in that we see a constant – although elevated 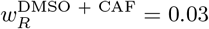 – growth rate in resistant cells; but a linearly decreasing (in *p*) growth rate of parental cells: 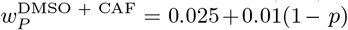 (or, equivalently: 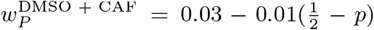). This could be interpreted as CAFs switching the direction of commensalism between parental and resistant cells.

### Leader and Deadlock games in NSCLC

The tools of evolutionary game theory (EGT) are well suited for making sense of frequency-dependent fitness [18–20, 26–30]. In EGT, a game is the rule mapping the population’s strategy distribution to the fitness of individual strategies. Previous work has considered games like snowdrift [24], stag hunt [25], rock-paper-scissors [21], and public goods [23, 26] alongside experiments. Instead, we experimentally operationalize the effective game (see [15, 16]) as an assayable hidden variable of a population and its environment. We define the effective game as the game played by an idealized population that shows the same frequency dynamics as the experimental population under consideration. As such, we are not aiming to test EGT as an explanation. Instead, we are defining a game assay to quantitatively describe our system in the language of EGT. In future work, it would be interesting to ask about the best language for describing cancer evolution by testing the game assay against several clearly and well operationalized alternatives to EGT.

To measure the effective game that describes the non-cell-autonomous interactions in NSCLC, we focus on the gain function (see [20, 31] for a theoretical perspective): the difference in growth rate between resistant and parental cells as a function of proportion of parental cells. The relatively good fit of a linear dependence of growth rates on parental seeding proportion allows us to describe the interaction as a matrix game – a well-studied class of evolutionary games (see a description in Figure 4a). Note that this linearity is not guaranteed to be a good description for arbitrary experimental systems. For example, the game between the two Betaproteobacteria *Curvibacter* sp. AEP1.3 and *Duganella* sp. C1.2 was described by a quadratic gain function [25]. If one views our work from the perspective of model selection then in the main text we proceed from the assumption of linearity. Supplemental Appendix F relaxes this assumption, extends our game assay to non-linear games, and compares linear and non-linear models with information criteria. Our qualitative results are unchanged, although the exact quantitative results for non-linear models differ slightly.

Two strategy matrix games have a convenient representation in a two dimensional game-space (see the model in Figure 4a and Supplemental Appendix C for details). This is the output of our game assay. We plot the inferred games in a game-space spanned by the theoretical fitness advantage a single resistant invader would have if introduced into a parental monotypic culture versus the fitness advantage of a parental invader in a resistant monotypic culture; as shown in Figure 4b. In this representation, there are four qualitatively different types of games corresponding to the four quadrants, each of which we illustrative with a dynamic flow. We can see that the game corresponding to DMSO + CAF – although quantitatively similar to DMSO – is of a qualitatively different type compared to all three of the other combinations.

We can also convert our inferred fitness functions from Figure 3 into a payoff matrix. We do this by having each row correspond to a strategy’s fitness function with the column entries as the *p* = 1 and *p* = 0 intersects of this line of best fit. These payoff matrix entries are abstract phenomenological quantities that could be implemented by various biological or physical processes [15]. If we look at our empirical measurements for DMSO + CAF (upper-right quadrant Figure 4b) we see the Leader game, and Deadlock in the other three cases (we will use DMSO to illustrate the Deadlock game).

**Figure 4:**
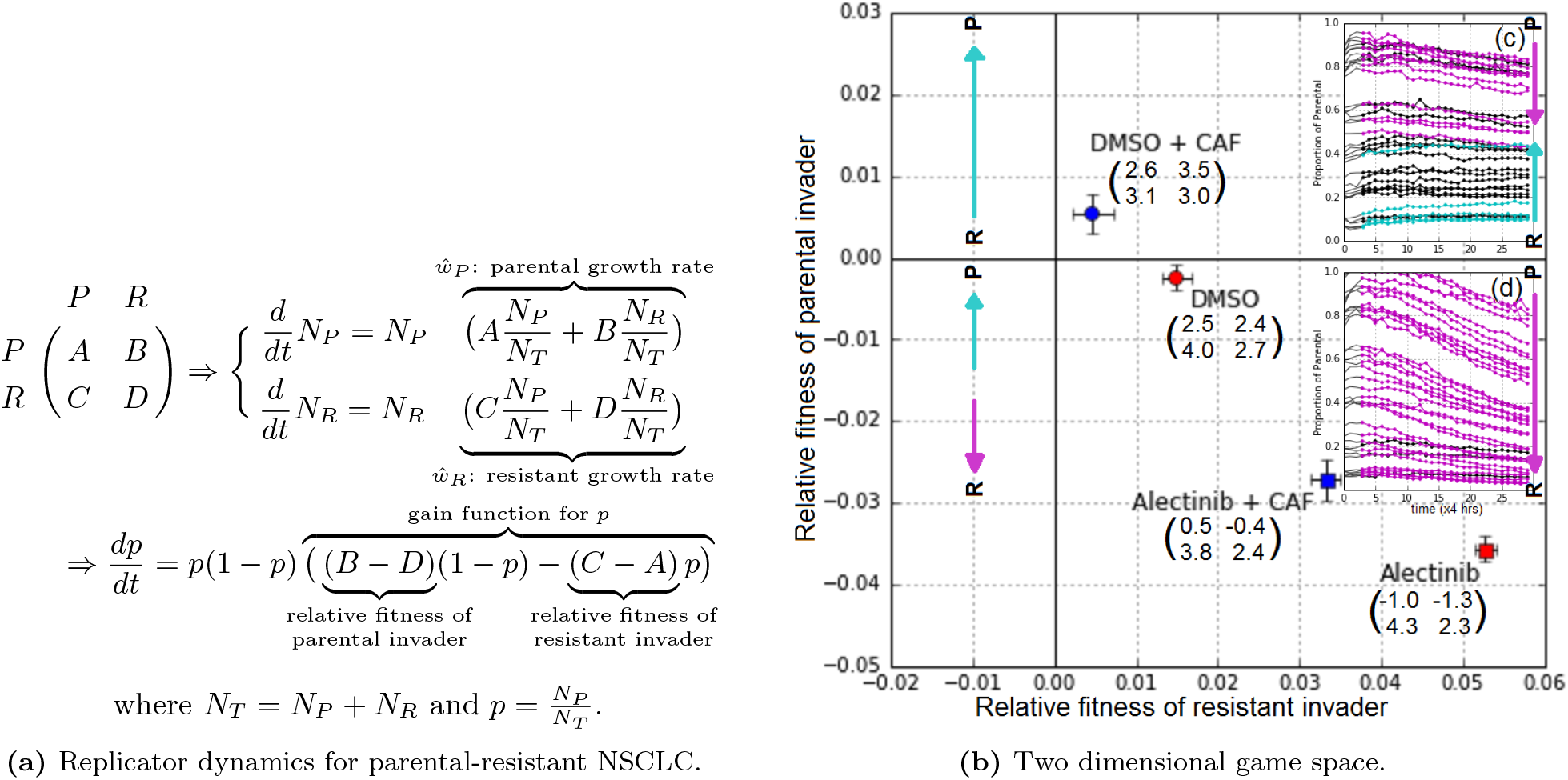
Measured games. (a) Replicator dynamics. Consider an idealized population of two strategies in a competitive co-culture: parental (P) and resistant (R). When a subpopulation of *P* interacts with *P* the subpopulation experiences a fitness effect A; when *P* interacts with *R* then *P* experience fitness effect *B* and *R* a fitness effect C; two Rs interact with fitness effects D, summarized in the matrix. This can be interpreted as an idealized exponential growth model for the number of parental (*N_P_*) and resistant (*N_R_*) cells. The dynamics of the proportion of parental cells 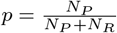 over time is described by the replicator equation (bottom). In Supplementary Section E we discuss a purely experimental interpretation based on Kaznatcheev [15]. **(b) Mapping of the four measured *in vitro* games into game space**. The x-axis is relative fitness of a resistant focal in a parental monotypic culture: *C* – *A*; y-axis is relative fitness of a parental focal in a resistant monotypic culture: *B* – *D*. Games measured in our experimental system are given as specific points with error bars based on goodness of fit of linear fitness functions in Figure 3. The games corresponding to our conditions are given as matrices (with entries multiplied by a factor of 100) by their label. See Supplemental Appendix C for more details. The game space is composed of four possible dynamical regimes, one for each quadrant. The typical dynamics of each dynamic regime are represented as qualitative flow diagram between *P* and *R*: an upward cyan arrow corresponds to an increase in the parental proportion, and a downward magenta arrow correspond to an increase in the resistant proportion. In the case of the two dynamic regimes observed in our NSCLC system, we also include insets of measured dynamics (c,d): **Experimental time-series of proportion of parental cells for DMSO + CAF (c) and Alectinib + CAF (d)**. Each line corresponds to the time dynamics of a separate well. A line is coloured magenta if proportion of resistant cells increased from start to end; cyan if proportion of parental cells increased; black if statistically indistinguishable proportions at start and end (where start/end are defined as the first/last 5 time-pints (20 hours)). See Supplementary Figure 5 for proportion dynamics of all four games and Supplementary Figure 6 for density dynamics and their correspondence to the exponential growth model from Figure 4a.

The Deadlock game observed in DMSO is in some ways the opposite of the popular Prisoner’s Dilemma (PD) game (in fact, Robinson & Goforth [32] call it the anti-PD). If we interpret parental as cooperate and resistant as defect then, similar to PD, each player wants to defect regardless of what the other player does (because 4.0 > 2.5 and 2.7 > 2.4; payoff numbers used in these examples are from the matrix entries we measured in Figure 4) but hopes that the other player will cooperate (because 4.0 > 2.7). However, unlike PD, mutual cooperation does not Pareto dominate mutual defection (because 2.5 < 2.7) but is instead strictly dominated by it. Thus, the players are locked into defection. In our system, this corresponds to resistant cells having an advantage over parental in DMSO.

The Leader game observed in DMSO + CAF is one of Rapoport [33]’s four archetypal 2 × 2 games and a social dilemma related to the popular game known as Hawk-Dove, Chicken, or Snowdrift (in fact, Robinson & Goforth [32] call it Benevolent Chicken). If we interpret parental as ‘lead’ (for Snowdrift: wait) and resistant as ‘work’ (for Snowdrift: shovel) then similar to Snowdrift, mutual work is better than both leading (because 3.0 > 2.6) and thus no work being done (for Snowdrift: both waiting and thus not getting out of the snowdrift) but each player would want to lead while the other works (because 3.5 > 3.0). However, unlike Snowdrift, mutual work is not better than the “sucker’s payoff” of working while the other player leads (because 3.1 > 3.0). Rapoport [33] sees this as a tension with a player switching from a “natural” point of mutual work to lead and thus benefit both players (3.5 > 3.0, 3.1 > 3.0), but if the second player also does the same and becomes a leader then all benefit disappears (because 2.6 is the smallest payoff). In our system, this corresponds to cells in the tumour experiencing selective pressure to lose some but not all of its resistance in DMSO + CAF.

Note that the above intuitive stories are meant as heuristics, and the effective games that we measure are summaries of population level properties [15, 16]: the population is the player and the two types of cancer cells are the strategies. This means that the matrix entries should not be interpreted as direct interactions between cells, but as general couplings between subpopulations corresponding to different strategies. The coupling term includes not only direct interactions, but also indirect effects due to spatial structure, diffusible goods, contact inhibition, etc. But this does not mean that an effective game is not interpretable. For example, the Deadlock game captures the phenomenon of the resistant population always being fitter than parental (for example, in DMSO). We noted this effect intuitively in Figure 2 (also see section Cost of resistance) from replicates being above the *y* = *x* diagonal. Measuring a Deadlock game for DMSO with confidence intervals that do not extend outside the bottom right quadrant of the game space in figure 4b allows us to show the statistical significance of our prior intuitive understanding. In other words, effective games allow us to quantify frequency-dependent differences in growth rates.

## Discussion

### Cost of resistance

The classic model of resistance posits that the resistant phenotype receives a benefit in drug (in our case: Alectinib or Alectinib + CAF) but is neutral, or even carries an inherent cost, in the absence of treatment (DMSO or DMSO + CAF). For example, experimentalists frequently regard resistance granting mutations as selectively neutral in the absence of drug, and the modeling community often goes further by considering explicit costs like up-regulating drug efflux pumps, investing in other defensive strategies, or lowering growth rate by switching to sub-optimal growth pathways [3, 34]. If we limited ourselves to the monotypic assays of Figure 1, then our observations would be consistent with this classic model of resistance. But in co-culture, we observed that resistant cells have higher fitness than parental cells in the same environment, even in the absence of drug. This is not consistent with the classic model of resistance. This higher fitness of resistant cells might not surprise clinicians as much as the biologists: in clinical experience, tumours that have acquired resistance are often more aggressive than before they were treated, even in the absence of drug. See Supplemental Appendix A.3 for a contrast of the biologist and clinician’s view of resistance in this context.

### Treating the game

Measuring a linear gain function has enabled us to develop an assay that represents the inter-dependence between parental and resistant cells as a matrix game. Experimentally cataloging these games allows us to support existing theoretical work in mathematical oncology that considers treatment (or other environmental differences) as changes between qualitatively different game regimes [18–20, 30]. In this framework, treatment has the goal not to directly target cells in the tumour, but instead to perturb the parameters of the game they are playing to allow evolution to steer the tumour towards a more desirable result (for examples, see [18–20, 30, 35, 36]). Empirically, this principle has inspired or built support for interventions like buffer therapy [37], vascular renormalization therapy [38], and adaptive therapy [39] that target the micro-environment and interactions instead of just attacking the cancer cell population. The success of the Zhang *et al*. [39] trial suggests that therapeutic strategies based on modulating competition dynamics are feasible. This highlights the need for a formal experimental method like our game assay that directly measures the games that cancer plays and tracks if and how they change due to treatment.

In our system, we can view an untreated tumour as similar to DMSO + CAF and thus following the Leader game. Treating with Alectinib (move to Alectinib + CAF) or eliminating CAFs through a stromal directed therapy (move to DMSO), moves the game into the lower-right quadrant of Figure 4b, and the game becomes a Deadlock game. Not only are these games quantitatively different among the four environmental conditions – see Figure 4b – but they are also of two qualitatively different types. To our knowledge, neither of the Leader and Deadlock games are considered in the prior EGT literature in oncology. Given that the Deadlock of drug-resistant over drug-sensitive cells is a challenge for classic models of resistance we would be particularly interested in theoretical models of resistance that produce the Deadlock game. In addition to challenging theorists by adding two new entries to the catalogue of games that cancers play, this switch allows us to show that the theoretical construct of EGT – that treatment can qualitatively change the type of game – has a direct experimental implementation. Unfortunately, neither of our *in vitro* games would lead to a therapeutically desirable outcome if they occurred in a patient.

### Heterogeneity and latent resistance

A particularly important difference between Leader and Deadlock dynamics is the existence of an internal fixed point in Leader but not in Deadlock. Fixed points are a property of equilibrium dynamics: in the most general case, even on very long timescales these fixed points might not be realized due to the evolutionary constraints of population size [40] or computation [41, 42]. Thus, it is important to check to what extent this qualitative difference can translate to a quantitative difference in finite time horizons. In our system, we can see a quantitative difference in the convergence towards the fixed point in the DMSO + CAF condition of Figure 4c, and no such convergence in the other three cases (Figure 4d for Alectinib + CAF; Supplementary Figure 5). Since the strength of selection (magnitude of the gain function) is small near a fixed point, the change in *p* also slows in the DMSO + CAF condition. We provide a more robust analysis of this in Supplemental Appendices C and F. It would be of interest for future work to study the long-term experimental stability of these fixed points.

Since the DMSO + CAF condition is our closest to an untreated patient, it might have important consequences for latent resistance. Many classical models of resistance assume a rare preexistent mutant taking over the population after the introduction of drug. In our experimental system, however, if the resistant strategy is preexistent then negative frequency dependent selection will push the population towards a stable polyclonal tumour of resistant and sensitive cells before the introduction of drug. This allows for much higher levels of preexisting heterogeneity in resistance than predicted by the classical picture. As such, we urge theorists to reconsider the assumption of the rare pre-existing resistant clone.

Of course, our results are for a single *in vitro* system. But if similar games occur in vivo and/or for other cancers, then such preexisting heterogeneity could be a possible *evolutionary mechanism* behind the speed and robustness of treatment resistance to targeted therapies in patients. This could help explain the ubiquity and speed of resistance that undermines our abilities to cure patients or control their disease in the long term. We will not know this unless we set out to quantify the non-cell autonomous processes in cancer. Building a catalogue of the games cancers play – by adopting our game assay in other cancers, and other experimental contexts – can help resolve this and other questions.

## Methods

### Cell lines

H3122 cell line was obtained from Dr. E. Haura (Moffitt Cancer Center). Cell line identity was validated by the Moffitt Cancer Center Molecular Genetics core facility using short tandem repeats (STR) analysis. Primary lung cancer associated fibroblasts were obtained from Dr. S. Antonia lab (Moffitt Cancer Center), following the protocols approved by the USF Institutional Review Board. CAFs were isolated as previously described in Mediavilla-Varela *et al*. [43] and expanded for 3-10 passages prior to the experiments. The Alectinib resistant derivative cell line was obtained through escalating inhibitor concentration protocol, as described in Dhawan *et al*. [13]. Alectinib sensitive parental H3122 cells were cultured in DMSO for the same length of time, as the alectinib resistant derivate.

Stable GFP and mCherry expressing derivative cell H3122 cell lines were obtained through lentiviral transduction with pLVX-AcGFP (Clontech) and mCherry (obtained from K. Mitsiades, DFCI) vectors, respectively. We cultured both H3122 cells and CAFs in RPMI media (Gibco brand from Thermo Scientific), supplemented with 10% FBS (purchased from Serum Source, Charlotte, NC). Regular tests for mycoplasma contamination were performed with MycoScope PCR based kit from GenLantis, San Diego, CA.

### Experimental set-up

The cells were harvested upon reaching 70% confluence and counted using Countess II automatic cell counter (Invitrogen). CAFs were counted manually to avoid segmentation artifacts. Mixtures of parental and resistant H3122 cells were prepared at 8 different ratios: all-resistant, 9:1 resistant to parental, 4:1, 3:2, 2:3, 1:4, 1:9, and all-parental. For the determination of competitive growth rates, 2,000 H3122 cells from the 8 mixtures were seeded with or without 500 CAF cells in 50 µL RPMI media per well into 384 well plates (Corning, catalogue #7200655), with different ratios of differentially labelled parental and alectinib resistant variants: with 6 wells used for each resistant:parental ratio in each of the 4 conditions. 20 hours after seeding, Alectinib – purchased from ChemieTek (Indianapolis, IN) – or DMSO vehicle control, diluted in 20 µL RPMI was added to each well, to achieve final Alectinib concentration of 500 nM/L [14]. Time lapse microscopy measurements were performed every 4 hours in phase-contrast white light, as well as green and red fluorescent channels using Incucyte Zoom system from Essen Bioscience.

### Game assay

We use the exponential growth rate in the fluorescent area of the two fluorescent channels as our measure of fitness. In order to minimise the impact of growth inhibition by confluency, we analyzed the competitive dynamics during the first 5 days of culture, when the cell population was expanding exponentially. We learned growth rate along with a confidence interval from the time-series of population size in each well using the Theil-Sen estimator. More detail on and justification of this measure of fitness is available in Supplemental Appendix B.2.

Since raw population sizes have different units (GFP Fluorescent Area (GFA) vs mCherry Fluorescent Area (RFA)), we converted them to common cell-number-units (CNU) by learning the linear transform that scales GFA and RFA into CNU. We defined proportions based on this common CNU as *p* = *N_P_/*(*N_P_ + N_R_*) where *N*{_*P,R*_} is the CNU size of parental and resistant populations. The transform of GFA to RFA into CNU is associated with an error that is propagated to measures of *p* as *σ_p_*.

To measure the fitness functions we plotted fitness of each cell-type in each well vs seeding proportion (*p*) of parental cells in Figure 3. The x-axis proportion of parental cells (*p*) was computed from the first time-point. We estimated the line of best-fit and error on parameters for this data using least-squares weighted by the inverse of the error on each data point. For the exact lines of best-fit, see Supplemental Appendix C.3.

The *p* = 0 and *p* = 1 intercepts of the lines of best fit serve as the entries of the game matrices. Note that in Figure 4b, we multiplied the entries by 100 for easier presentation. The game point are calculated from the matrices as *x* := *C — A* and *y* := *B — D*, and the error is propagated from the error estimates on lines of best-fit’s parameters.

### Data Availability

Due to size constraints, raw image data from experiments are available upon request. Post-image processing data (i.e. population size time-series for each experimental replicate) is available on GitHub at https://github.com/kaznatcheev/GameAssay.

### Code Availability

Image analysis code is available on GitHub at https://github.com/kaznatcheev/CV4Microscopy. The game assay analysis code is available on GitHub at https://github.com/kaznatcheev/GameAssay.

### Contributions

AK, JP, AM, and JGS conceived and designed the study. JP and AM performed the experiments. AK designed the mathematical model, wrote the image analysis and game assay code, and analysed the data. AK, AM and JGS wrote the main text. AK and JP wrote the supplemental appendices. DB, AM, and JGS supervised the project. All authors discussed the results and implications; commented on the work at all stages; and approved the final submission.

## Acknowledgements

JGS would like to acknowledge the NIH Loan Repayment program for their generous support of his research in general as well as Miles for Moffitt and NIH Case Comprehensive Cancer Center support grant P30CA043703 and the Calabresi Clinical Oncology Research Program, National Cancer Institute of Award Number K12CA076917. We would also like to thank Mohamed Abazeed, Peter Jeavons, Konstantine Kaznatcheev, and three anonymous reviewers for helpful feedback and discussions.

## Appendices

In these supplementary appendices, we develop and discuss the tools used to define and measure our game assay. The structure is as follows:

A. Description of materials used, the experimental method, and time-lapse microscopy. Discussion of resistance terminology.
B. Basic quantification of experimental images and how growth rates and associated error are measured within each well. This is the definition of fitness used throughout our text. Explains Figures 1 and 2 from the main text.
C. Defines parental proportions (*p*) and contrasts the evolutionary dynamics of proportions (Supplementary Figure 5) with the ecological dynamics of densities (Supplementary Figure 6). Defines the fitness functions based on lines of best between fitness and proportion from Figure 3 and presents the actual lines of best fit as equations 1-8. Explains how the linear fitness functions are converted into gain functions and games; and how those games are plotted in Figure 4b. Justifies the use of linear functions in terms of explanatory value and presents the model residuals in Supplementary Figure 7.
D. Presentation of interpretable fitness functions 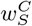 from section ‘Frequency dependence in fitness functions’ of the main text in the context of regularization. As a visual check of the regularization, Supplementary Figure 8 shows what the games would look like if based on the regularized fitness functions 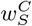 instead of the unregularized fitness functions 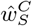 shown in Figure 4b.
E. Experimental interpretation of replicator dynamics as an alternative to exponential model of Figure 4a.
F. Generalization of game assay to non-linear fitness functions. Provides the 3rd order fitness functions (Supplemental Figure 9) or independent mixed order fitness functions (Supplemental Figure 11) that would be selected by information criteria (Supplemental Figure 10) if one treats the game assay not as a definition of games but as a model selection problem for parameter fitting. As a qualitative check, Supplementary Figure 12 shows the agreement in game space between the higher order games and our measured matrix games.

### A Materials and experimental method

#### A.1 Cell lines

H3122 cell line was obtained from Dr. E. Haura (Moffitt Cancer Center). Cell line identity was validated by the Moffitt Cancer Center Molecular Genetics core facility using short tandem repeats (STR) analysis. Primary lung cancer associated fibroblasts were obtained from Dr. S. Antonia lab (Moffitt Cancer Center), following the protocols approved by the USF Institutional Review Board. CAFs were isolated as previously described in Mediavilla-Varela *et al*. [43] and expanded for 3-10 passages prior to the experiments. The alectinib resistant derivative cell line was obtained through escalating inhibitor concentration protocol, as described in Dhawan *et al*. [13]. Alectinib sensitive parental H3122 cells were cultured in DMSO for the same length of time, as the alectinib resistant derivate.

Stable GFP and mCherry expressing derivative cell H3122 cell lines were obtained through lentiviral transduction with pLVX-AcGFP (Clontech) and mCherry (obtained from K. Mitsiades, DFCI) vectors, respectively. We cultured both H3122 cells and CAFs in RPMI media (Gibco brand from Thermo Scientific), supplemented with 10% FBS (purchased from Serum Source, Charlotte, NC). Regular tests for mycoplasma contamination were performed with MycoScope PCR based kit from GenLantis, San Diego, CA.

#### A.2 Experimental set-up

The cells were harvested upon reaching 70% confluence and counted using Countess II automatic cell counter (Invitrogen). CAFs were counted manually to avoid segmentation artifacts. Mixtures of parental and resistant H3122 cells were prepared at 8 different ratios: all-resistant, 9:1 resistant to parental, 4:1, 3:2, 2:3, 1:4, 1:9, and all-parental. For the determination of competitive growth rates, 2,000 H3122 cells from the 8 mixtures were seeded with or without 500 CAF cells in 50 µ*L* RPMI media per well into 384 well plates (Corning, catalogue #7200655), with different ratios of differentially labelled parental and alectinib resistant variants: with 6 wells used for each resistant:parental ratio in each of the 4 conditions. 20 hours after seeding, Alectinib – purchased from ChemieTek (Indianapolis, IN) – or DMSO vehicle control, diluted in 20 µL RPMI was added to each well, to achieve final Alectinib concentration of 500 nM/L [14]. Time lapse microscopy measurements were performed every 4 hours in phase-contrast white light, as well as green and red fluorescent channels using Incucyte Zoom system from Essen Bioscience.

#### A.3 Reductive vs effective definitions of resistance

In these experiments, we observed (see Figure 2 and Sections ‘Monotypic vs mixed cultures’ and ‘Cost of resistance’) that even in the absence of drug, resistant cells tend to have a higher growth rate than parental cells in the same environment (i.e. proportion of parental cells in the co-culture). A reductionist could rationalize our observations by saying that we actually selected for two different qualities in our resistant line: (i) a general growth advantage, and (ii) resistance to Alectinib.

This is a reasonable hypothesis, but it faces a few challenges. First, both parental and resistant cells were evolved for the same length of time, with escalating dosages of DMSO for the former and Alectinib for the latter (see Mediavilla-Varela *et al*. [43] and above). Thus, (i) cannot be due to just subculturing, but is somehow linked to drug. Second, there is no growth rate advantage of resistant cells in monoculture (see Figure 1); the advantage is only revealed when parental and resistant cells are cultured with a common proportion of parental cells. Finally, to even make the distinction between (i) and (ii), one has to implicitly assume that resistance has to be neutral or costly by definition. For an oncologist, however, both (i) and (ii) would constitute clinical resistance if they led to a tumour escaping therapeutic control. By using a definition of clinical resistance that is broad enough to capture both aspects, we observe resistance that is neither neutral nor costly in DMSO co-culture.

### B Measuring population sizes and fitnesses

#### B.1 Fluorescent area as units of population size

We measured fluorescent area from time-lapse images via python code using the OpenCV package and used this as our units of size for populations. See Kaznatcheev [15] for a discussion of fitness and replicator dynamics under various definition of population size. We cleaned images by renormalizing them (GFP and mCherry intensities vary over different orders of magnitude), removed vignetting with CLAHE, and finally thresholded to identify fluorescent regions. We eliminated salt-and-pepper noise from the thresholded images with the opening morphological transform. See Figures 2a,b,d,e for examples of the image analysis. The resultant area is then taken as a measure of population size for the purposes of computing fitnesses.

#### B.2 Growth rate as fitness

We use the exponential growth rate (or Malthusian parameter) as our measure of fitness. In order to minimise the impact of growth inhibition by confluency, we analyzed the competitive dynamics during the first 5 days of culture, when the cell population was expanding exponentially. See section E for a discussion of the impact of measurement length. We learned growth rate along with a confidence interval from the time-series of population size in each well using the Theil-Sen estimator [44, 45]. Since the Theil-Sen estimator is a rank-based median method (unlike least-squares, which is a numeric-based mean method), it is more robust to noise and does not need to choose between a linear or log representation for computing the error-term (since log transforms do not change rank orders). The robustness to rare but large magnitude noise is useful for our purposes because such errors do not reflect biological function or noise but are more likely to be due to errors in image processing, for example in response to sudden condensation on the well plate. See Figures 2c,f for examples of fitting.

The learned parental growth rate and resistant growth rate of each well are used as the *y* coordinates in the monoculture experiments of Figure 1 (along with errors on the growth rate) and as the *x* and *y* coordinates of the main part of Figure 2. Due to too much information content, the errors on the growth rates are omitted in Figure 2, but they are shown explicitly as error-bars in Figure 3. Note that this means that each point in Figure 1 and the main part of Figure 2 and each pair of points in Figure 3 (one magenta and one cyan at the same x-position) correspond to one biological replicate, with the error term coming from the confidence interval on the growth-rate estimate from the 30 time-series points that we recorded for each biological replicate (see section E on how this relates to the accuracy-precision trade-off). Thus, each of the 6 wells corresponding to a given resistant:parental ratio (in each of the 4 conditions) has its own independent growth rate with associated error. The wells are not averaged together: each acts as its own data point (with noise) for later analysis (that propagates the noise).

#### B.3 Other definitions of fitness

Of course, in the most general case, it is possible to consider other alternatives to the exponential growth rate or Malthusian parameter as definitions of fitness. Popular alternatives include logistic growth rate and more general Gompertz growth rate, but many choices are possible. However, there are experimental, conceptual, and mathematical reasons for why we focus on the exponential growth model.

Experimentally: if the exponential growth model is a poor choice for our game assay pipeline, this will show up in an unreasonably large error term on the growth rate, which would then propagate to inconclusive measurements of the game. This is one of the advantages of being able to estimate error terms for the growth rate of each individual biological replicate (instead of the more common practice of relying on variance between replicates). In other words, if the error bars are big on the growth rate, then this will increase the size of error bars on the final game measurement – potentially to the point that the game cannot be localized with confidence to a given quadrant of game space. For our particular experimental system, this is not the case, and the error terms on growth rate are sufficiently small to get conclusive measurements of the game.

Conceptually: an effective game is defined with respect to a choice of idealized population. It is better if the fitness measure is natural for that idealized population. In our case, both the intuitive presentation of games in Figure 4a and operational presentation in section E are purely multiplicative models. And exponential growth is the generator of multiplicative models.

Mathematically: a central advantage of representing evolutionary dynamics as games is to make qualitative distinctions between types of games. The qualitative nature of a game depends only on the rank ordering of its payoff matrix entries. Any strictly monotonic transform between fitness value will not change the rank ordering of payoff matrix entries and thus preserve the main qualitative conclusions of a game theoretic analysis. Thus, in the context of the game assay of our experimental system, little is to be gained from a more complex definition of fitness.

#### B.4 Figure 2 as map of analysis flow

Along with showing all the data, Figure 2 serves as a map to the above analysis pipeline. The subfigures can be understood in the following order:

[a,b,c,d]: Within each image from the series generated by time-lapse microscopy: identify the fluorescent regions for GFP and mCherry and calculate their areas to serve as units of population size (GFA and CFA).
[e,f]: For parental (mCherry) and resistant (GFP) plot the population sizes from each image in the series on a semilog grid as population vs. time. Find the slope of the two lines to serve as parental and resistant fitness.
[g]: Use the parental fitness as *x* value and resistant as *y* value to plot each well as a data-point according to the above process, and color the point according to its experimental condition (with opacity for initial parental proportion; see Section C). For ease of viewing: put a convex hull binding polygon around each well data-point dependent on their experimental condition.

Given the complexity of Figure 2, it is tempting to ask for a simple summary statistic of the data in the main figure. But it is not reasonable to ask of the “average” growth rate in Figure 2 because each point differs not only along the four experimental conditions of the environment, but also along the micro-environmental conditions of the initial parental proportion (represented by the opacity that is explained in SA C). Averaging over this information would be akin to assuming that the growth rates are cell-autonomous. It would be attributing the variance in growth rates to noise instead of the independent variable or initial parental proportion. As such, the game assay developed in rest of the paper can be viewed as a method for summarizing Figure 2 when the underlying process is non-cell-autonomous. And the games derived through Figure 3 and presented in Figure 4b are the summary of the data in the main part of Figure 2.

**Supplementary Figure 5:**
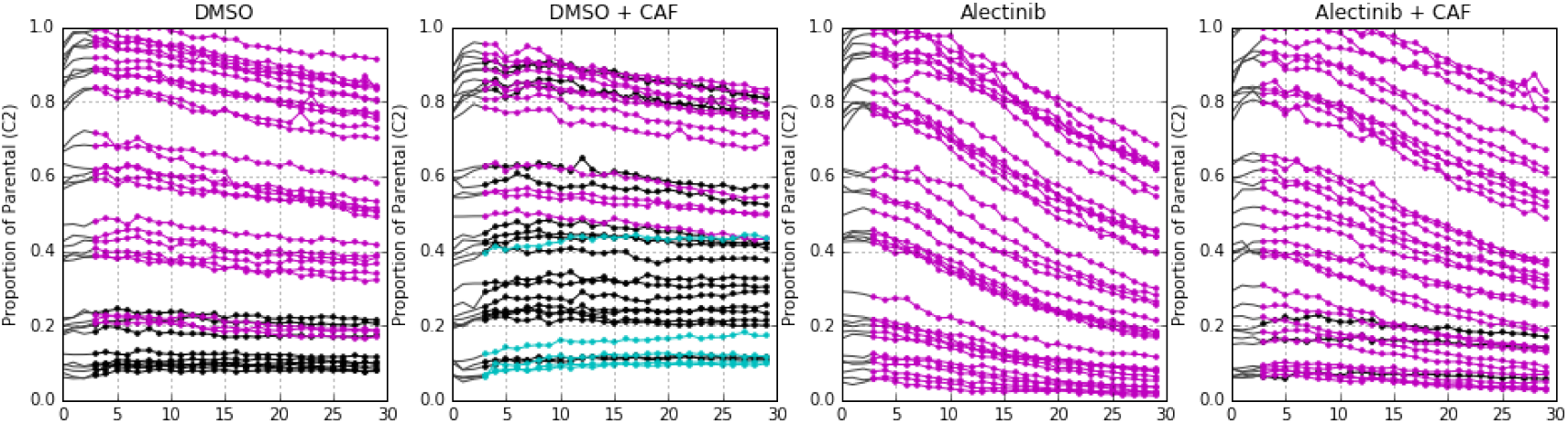
Evolutionary dynamics of proportion of parental cells versus time for competition of parental vs. resistant NSCLC. Each line corresponds to the time dynamics of a separate well. A line is coloured magenta if proportion of resistant cells increased from start (time step 3 to 8) to end (time step 24 to 29); cyan if proportion of parental cells increased; black if statistically indistinguishable proportions at start and end.

### C Measuring fitness functions and games

#### C.1 Proportions

Since raw population sizes have different units (GFP Fluorescent Area (GFA) vs mCherry Fluorescent Area (RFA)), we converted them to common cell-number-units (CNU) by learning the linear transform that scales GFA and RFA into CNU. We defined proportions based on this common CNU as *p* = *N_P_*/(*N_P_ + N_R_*) where *N*{_*P,R*_} is the CNU size of parental and resistant populations. The transform of GFA to RFA into CNU is associated with an error that is propagated to measures of *p* as *σ_p_*. Thus, although we used 8 different ratios of resistant to parental cells with 6 wells per condition seeded at each of the ratios, we do not average over these 6 wells but associated each with its own proportion *p* ± σ_p_ from the initial image. This helps us control for systemic noise from field of view and our image processing algorithm. The time dynamics of *p* can be seen in the insets of Figure 4b for DMSO and DMSO+CAF or in Supplementary Figure 5 for all conditions.

#### C.2 Neglecting ecological dynamics

Throughout this report, we focus on evolutionary dynamics: changes in proportion of strategies. However, one could also consider the ecological dynamics: changes in densities of strategies. It is not only proportions that are changing in our experimental system but also the densities. These ecological dynamics are not the focus of our report, but we present them in Supplementary Figure 6 for completeness. Here, we also compare the prediction of the model based on our measured games and the exponential growth interpretation in Figure 4a to the observed data. There is overall agreement between data and model. But this is based on the traditional two track approach on qualitative agreement. Instead, we prefer to focus on the single track measurement of evolutionary dynamics described in the rest of these supplementary materials. Future work can aim to extend our approach to also include ecological dynamics.

#### C.3 Lines of best fit as fitness functions

To measure the fitness functions we plotted fitness of each cell-type in each well vs seeding proportion (*p*) of parental cells in Figure 3. The x-axis proportion of parental cells (*p*) was computed from the first time-point: see section E for an interpretation of this as a measurement of *dp/dt* or as a series of competitive fitness assays. We estimated the line of best-fit and error on parameters for this data using least-squares weighted by the inverse of the error on each data point (i.e. 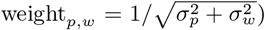. This provides the error estimates on the line’s parameters that we use later. The lines of best fit (with coefficients rounded to the thousandths for presentation) from weighted least-squares are:

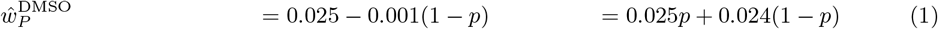

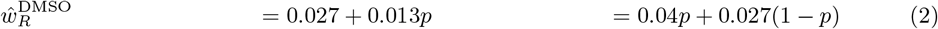

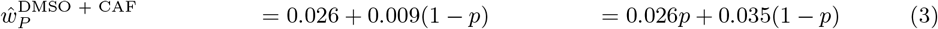

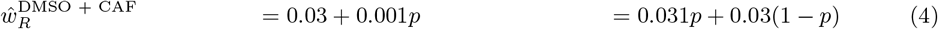

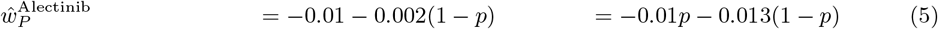

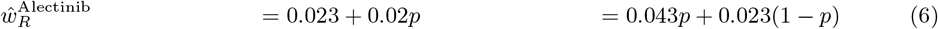

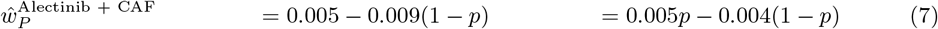

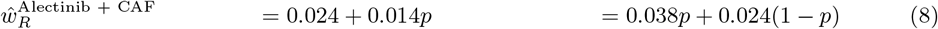

**Supplementary Figure 6:**
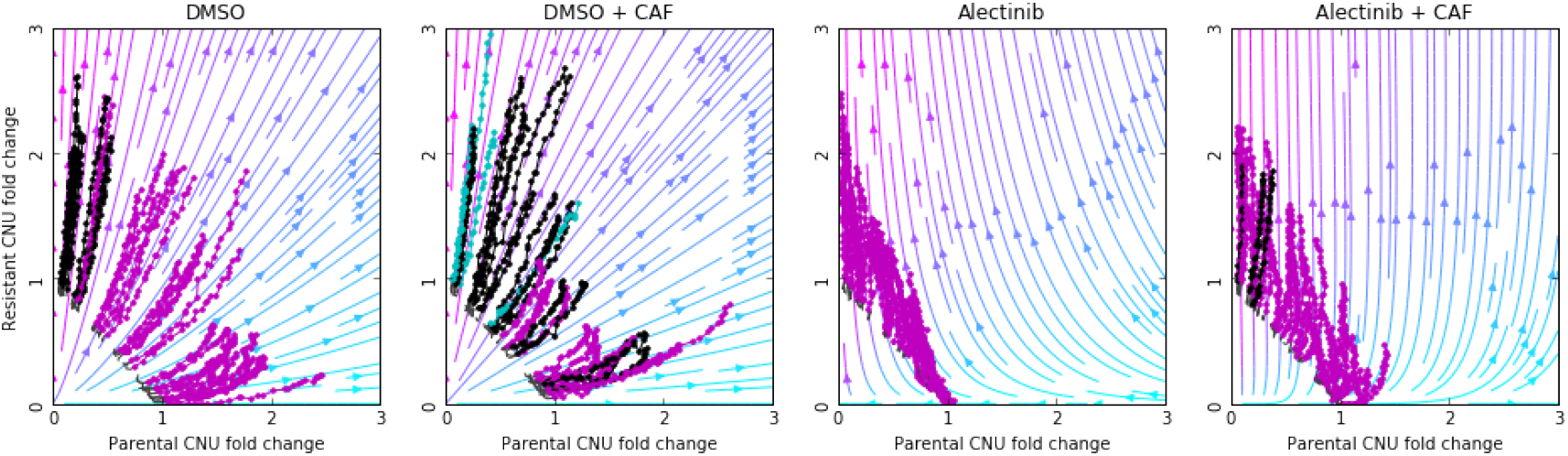
Dynamics of population sizes of resistant cells versus parental cells. Axis are fold change in CNU normalized from the seeding proportion of each well, with x-axis for parental and y-axis for resistant. Foreground: raw data. Each line corresponds to the time dynamics of a separate well. A line is coloured magenta if proportion of resistant cells increased from start to end; cyan if proportion of parental cells increased; black if statistically indistinguishable proportions at start and end (using the same conventions as Figure 5). Background: flow diagram for model from Figure 4a. Each coloured point shows the proportion of parental-resistant (cyan-magenta) at that point. Arrow going from magenta to cyan indicates parental proportion increased, if from cyan to magenta then resistant proportion increased.

#### C.4 Summarizing fitness functions as games

For the final column of our presentation of 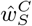 in equations 1-8, we rewrote the fitness functions in a suggestive form of 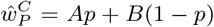 and 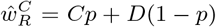. This is done to show at a glance where the matrix entries in Figure 4b come from. This is because the *p* = 0 and *p* = 1 intercepts of the fitness functions serve as the entries of the game matrices. Note that in Figure 4b, we multiplied the entries by 100 for easier presentation. The game point are calculated from the matrices as *x* := *C* – *A* and *y* := *B* – *D*, and the error is propagated from the error estimates on fitness function’s parameters.

#### C.5 Gain functions, game space, and fixed points

A particularly important equation for studying two strategy games is the gain function. This represents the relative fitness difference between two strategies. Thus, it is a measure of selection strength and a proxy for the rate of evolution. The parental gain function (i.e. gain function for *p* in Figure 4a and equation 23) is given by 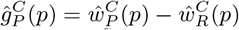; and the resistant gain function (i.e. gain function for 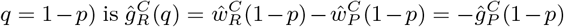. The end-points of this gain function determine the game coordinates in the game space of Figure 4b with 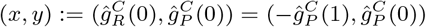. These points can be interpreted as the idealized quantities of relative fitness of a resistant invader in parental monoculture 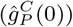 and relative fitness of a parental invader in resistant monoculture 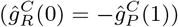. If these two coordinates have the same sign then the gain function has to cross 0 in getting from *p* = 0 to *p* = 1 and thus the dynamics have a fixed point. If the two coordinates are both positive (top right quadrant of Figure 4b) then the fixed point is stable, if both are negative (bottom left quadrant of Figure 4b) then the fixed point is unstable. In our experimental system, only the DMSO + CAF condition has a fixed point at 0.53 ± 0.14 (rounded to the nearest percent).

**Supplementary Figure 7:**
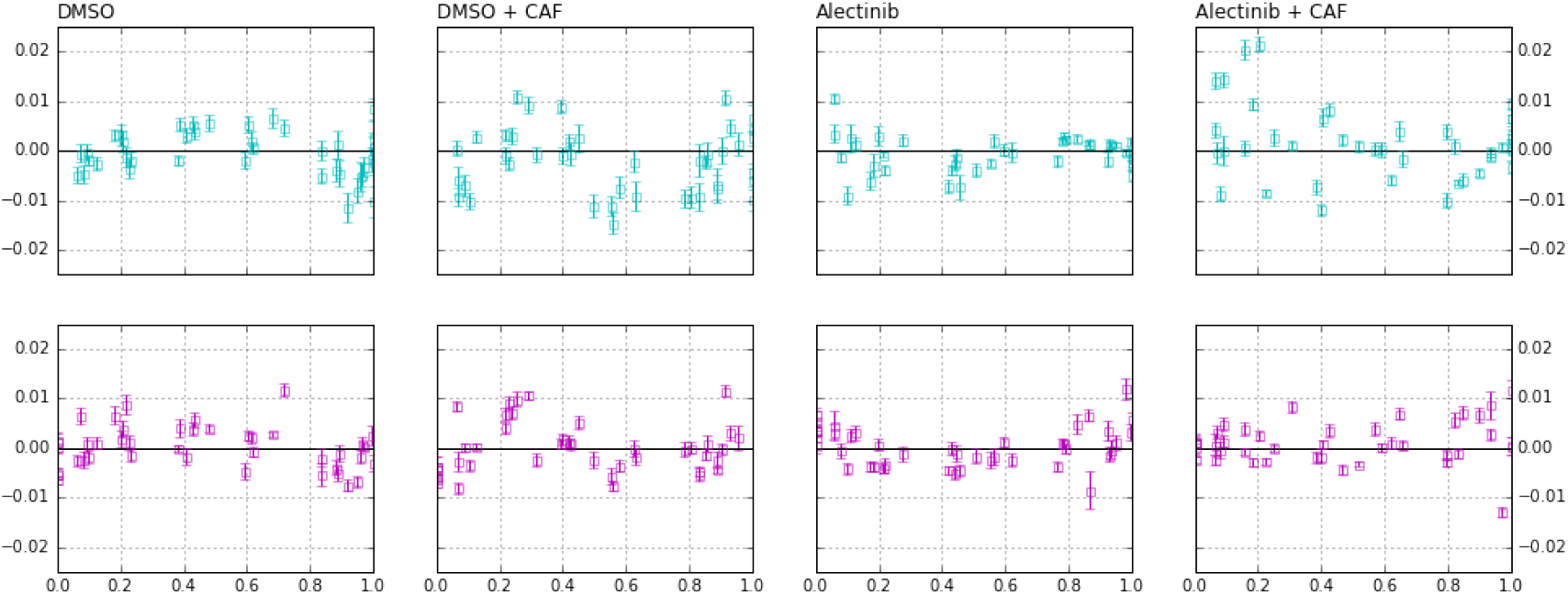
Residuals for the fitness functions. The x-axis is proportion and y-axis is residuals of the lines of best fit from Figure 3 for parental (Cyan,Top) and resistant (Magenta,Bottom).

#### C.6 Width and height of fixed regions

Since we propagate the errors on our measurement from the image all the way to the game, we find it more helpful to think of an experimental fixed point not as a point but as a fixed region *p* ∈ (0.39, 0.67) of finite width. This can provide an alternative explanation for the apparent slowness of convergence to the fixed point in Figure 5. Some of the fixed region’s width is noise in measurement, but some could be due to true variance between wells: in particular, even if the reductive game is the same, the spatial structure will be slightly different in each well and thus there will be a slightly different effective game. As such, apparent slowness in Figure 5 might be from different lines being very close to slightly different fixed points that are all within the fixed region’s width. An alternative view to width is in terms of height: a fixed region corresponds not just to the point where 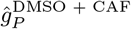 crosses 0 but to the region where the gain function crosses 0 ± 0.0014 (rounded to the nearest thousandth). We call this the fixed region’s height (and use it in section F). This height is due to propagation of error and can be interpreted as our measurement not being able to distinguish relative growth rates in (−0.0014,0.0014) from zero. In the case of the other three conditions (DMSO, Alectinib, and Alectinib + CAF), in going from *p* = 0 to *p* = 1, the gain function do not pass within their fixed region height of zero, and thus no fixed regions exist.

#### C.7 Lines and matrix games

Although slight deviations from a linear fit – that might not be attributable to noise alone – might be present in the data (see Supplementary Figure 7), we do not think that they justify considering higher-order fitness functions (although we discuss higher-order functions in section F for completeness). This is due to the higher explanatory value of linear models and our hope to influence the well-established study of matrix games in microscopic systems. Some good EGT work has recently been done on non-linear games [25, 26, 30], but this is very little compared to the immense literature on matrix games. More importantly, we think that our focus on matrix games is better viewed not from the perspective of model selection but rather as an operational definition of effective games. We are not aiming to provide the best or most predictive account of non-small-cell lung cancer in the petri dish, but rather a method for measuring (matrix) games. If the error of the measured (matrix) games ends up very high – which is not the case from the error bars in Figure 4b – then we know that this first order approximation of interactions is not sufficient and higher orders should be pursued. However, we will not know this unless we first have a robust method for measuring the lower order terms.

### D Regularization and interpretable fitness functions

Regularization is a machine learning technique for reducing over-fitting by biasing towards more succinct models. It is the use of *a priori* knowledge on what constitutes a simpler or more likely model to anchor our inference. A classic example of this is preferring lower-order over higher-order polynomials for describing data unless there is overwhelming evidence otherwise. Of course, what constitutes overwhelming evidence depends on the goals of the scientists. If the only goal is prediction then cross-validation is a good way to test how heavily inference should be regularized. But if the goal is explanation then accordance with existing theory is another important factor to consider.

**Supplementary Figure 8:**
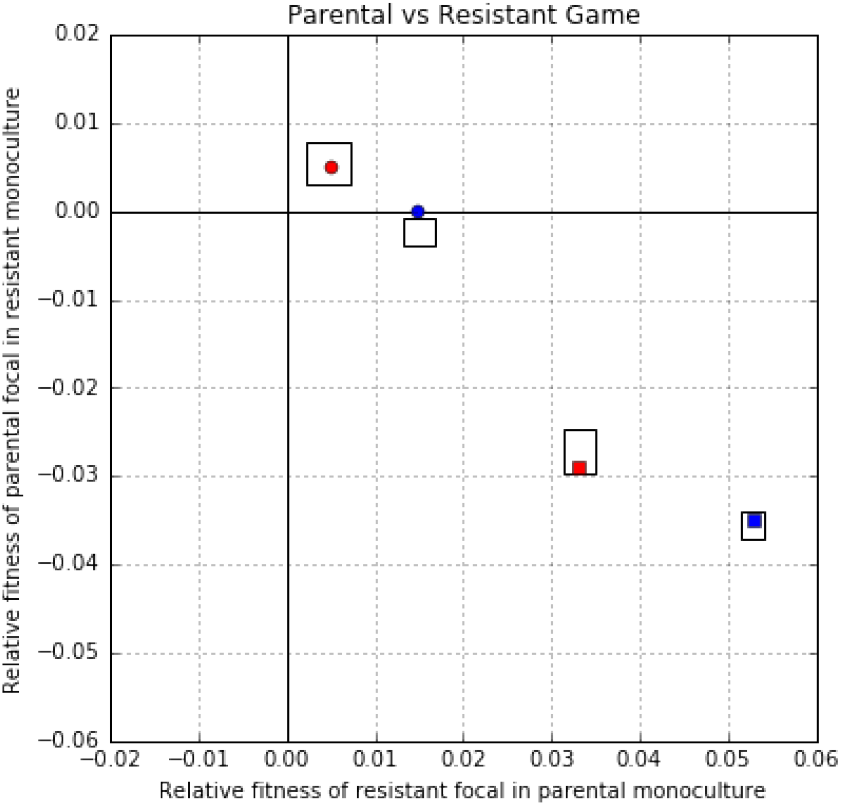
Mapping of the regularized fitness functions for the four conditions into game space. The x-axis is the relative fitness of a resistant focal in a parental monotypic culture: **C – A**. The y-axis is the relative fitness of a parental focal in a resistant monotypic culture: **B – D**. Games measured in our experimental system are specified by the bounding boxes corresponding to the range of their errors. The games corresponding to the regularized fitness functions in equations 9-16 are given as points. Experimental condition is represented by shape (DMSO: circle; Alectinib: square) and colour (no CAF: red; + CAF: blue).

As such, our choice of focusing on linear fitness function in section C and Figure 3 can be seen as a form of regularization. In particular, we can see our inference procedure as either restricted to the hypothesis class of linear functions, or as considering the hypothesis class of all polynomials but with prohibitively high costs for non-zero components (*l*_0_ regularization) on orders beyond linear. But we prefer to think of it in terms of operationalization. By introducing a game assay, we are defining the hidden variable of (matrix) games in terms of the measurement procedure that we described in sections B and C.

#### D.1 Interpretable fitness functions

An uncontroversial case of regularization in our report is the presentation of *w_S_^C^* in section ‘Frequency dependence in fitness functions’. There, we restrict beyond linear fitness functions to focus on conceptually simple ones. In particular, we favor cell-autonomous functions over frequency dependent ones (i.e. *l*_0_ regularization on the fitness function coefficients) and we favor coefficients that are shared between different *S* and *C*. This results in the following regularized fitness functions:

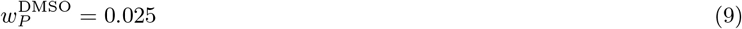

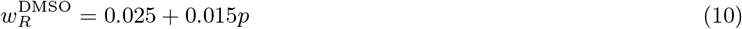

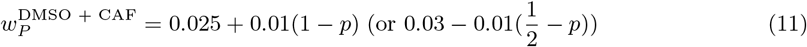

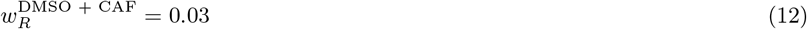

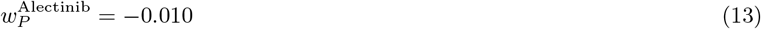

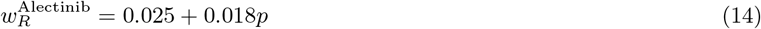

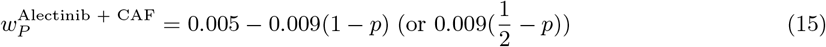

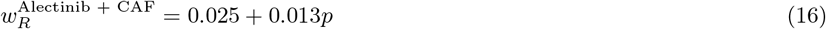

Note that for both *P* and *R* strategies, we used the proportion of the other strategy (1 – *p*, *p*) as the parameter that captures the non-cell-autonomous contribution. In equations 11,15, we also consider the parameter 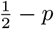 because of the elegant form it provides.

We can compare these regularized fitness functions 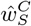 to the non-regularized 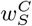 in equations 1-8. As can be seen, all 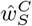 are close to their respective 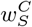 and are actually within the error estimates on 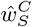. We can see the regularization in action with a push towards a constant base fitness of 0.025 shared by 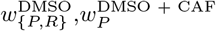, and 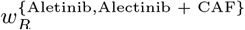. The absence of frequency dependent perturbation terms for 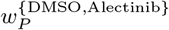 and 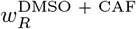 suggests that these strategies can be explained in terms of cell-autonomous processes. However, the other strategies in the other contexts ask for a non-cell-autonomous explanation.

#### D.2 Games from interpretable fitness functions

For a visual confirmation that the regularization of 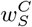 in equations 9-16 are reasonable, we can transform them into regularized games. We do this in the same way as we did for transforming the non-regularized 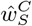 in equations 1-8 into the game-points of figure 4b. The results are in Supplementary Figure 8. The regularized games (points) are within the confidence rectangles of the measured games (boxes), with the exception of DMSO which is just outside its box. This is reasonable given that the boxes correspond to error: i.e. around 2/3rds confidence.

### E Experimental definition of replicator dynamics

Consider a well that is seeded with an initial number 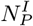 of parental and 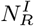 of resistance cells; total number 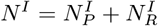. Let 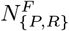 be the number of {parental,resistant} cells after being grown for an amount of time ∆*t*. From this, the experimental growth rate can be defined based on fold change as:

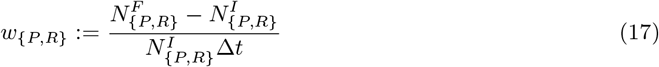

this can be rotated into a mapping *N^I^* → *N^F^* given by 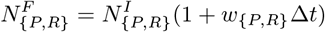.

By defining the initial and final proportion of parental cells as 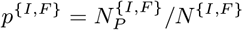, we can find the mapping:

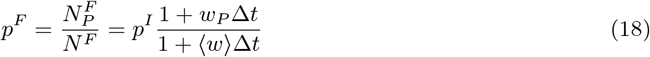

where 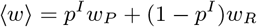. This is the discrete-time replicator equation.

We can approximate this discrete process with a continuous one by defining 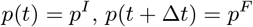 and looking at the limit as *∆*t** gets very small:

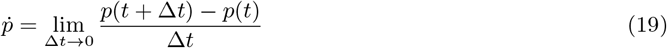

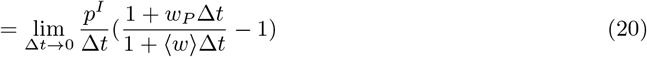

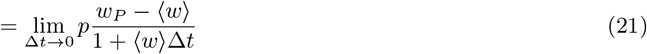

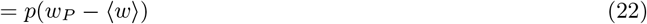

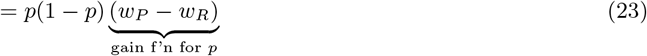

Thus, we recover replicator dynamics as an explicit experimental interpretation for all of our theoretical terms. Note that we did not make any assumptions about if things are inviscid or spatial; if we are talking about individual or inclusive fitness; or, if we have growing populations in log phase or static populations with replacement. All of these microdynamical details are buried in the definition of experimental fitness. This allows us to focus on effective games [15] and avoid potential confusions over aspects like spatial structure [16].

#### E.1 Better estimates of *w*

The problem with the definition of *w* in equation 17 is that it depends on just two time points, and thus not good for quantifying error. In our experimental system, we are able to peek inside the system with time-lapse microscopy. This allows us to get more than just the initial and final population sizes and replace fold-change by the more specific measurements of inferred growth rates for *w*_{*P,R*}_ that we describe in section B. An advantage of this approach is that the goodness-of-fit of the exponential growth model provides a good estimate of the error associated with each measurement of w. Thus, we are able to quantify error within each well and not just between experimental replicates in different wells with similar initial conditions.

**Supplementary Figure 9:**
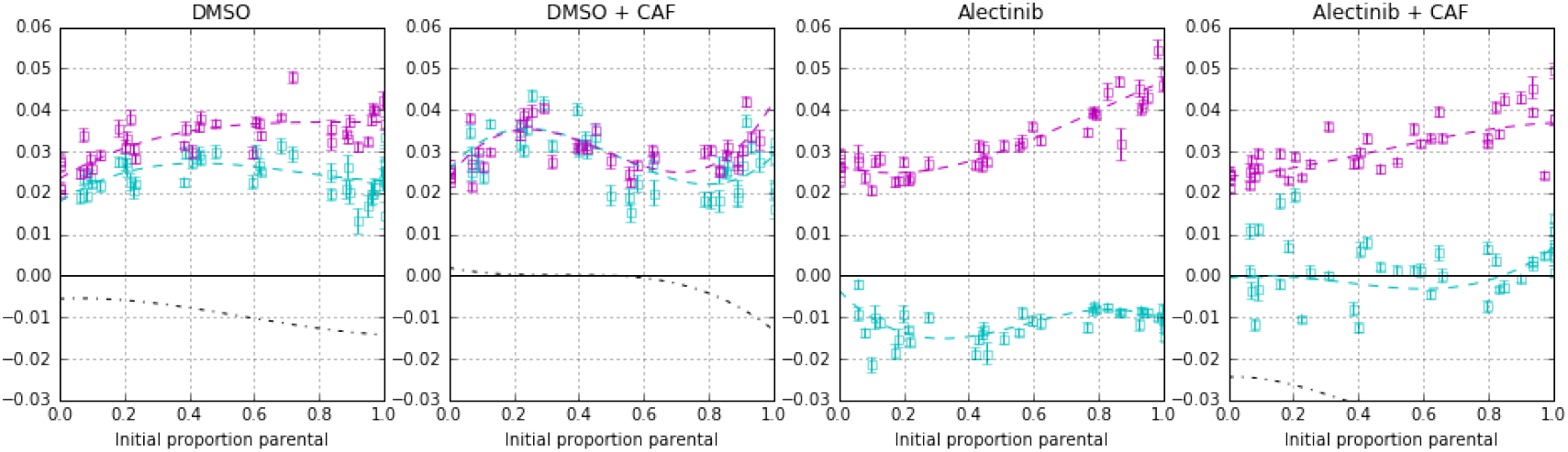
Cubic Fitness functions for competition of parental vs. resistant NSCLC. For each plot: growth rate with confidence intervals versus initial proportion of parental cells. This is the same data as Figure 3. Cyan data points are growth rates of parental cells, and magenta for resistant cells. Dotted lines represent the 3rd-order (cubic) fitness function of the least-squares best fit. The black dotted line is the gain function for parental (see Figure 4a), it is well below the *y* = 0 line in the Alectinib conditions (indicating the strong advantage of resistance) and thus cut out of the figure.

#### E.2 Accounting for finite ∆*t*

The small time definition of the derivative can be thought of as a way to approximate a function by local linearizations. It is why for simulations, modelers often use the discrete time replicator dynamics to represent continuous time replicator dynamics: effectively using the discretization as a simple ODE solver/plotter. In the limit of ∆*t* going to 0, this linearization recapitulates the function. Unfortunately in practice, our experimental system cannot take the limit as ∆*t* goes to 0 because of a precision-accuracy trade-off. Accuracy increases as ∆*t* decreases because the continuous dynamics is approximated by more and more, shorter and shorter straight lines. But – from an experimentalist’s perspective – the precision decreases because any measurement is noisy: if we measure growth rate over a shorter period of time then we are less certain whether our measurement reflects reality or noise. For very short measurements, we might get higher accuracy (assuming biological factors like time from seeding to adherence could be ignored) but would have incredibly low precision (due to only one, two or three time points from which to calculate growth rate). As we increase the time of the experiment, the accuracy might decrease but the precision will tend to increase. This is a classic trade off between random noise (low precision) and systematic noise (linearization being progressively less accurate over larger ∆*t*). Since each of our growth rate measurements has an associated error term (see section B), we quantify the random and systematic noise together and propogate it throughout our analysis. Given the biological constraints of our system, we judged that 5 days was a good trade-off point. This will most likely be different for other experimental systems.

Given that *w* are defined over a finite range of time, we need to pick a particular time-point to associate each measurement with. As is common for discrete time process, we attribute the value of the growth rate to the initial point. In particular, this means that when we make *w_{P,R}_* a function of *p* in the main text, then the values of growth rate are attributed to the initial proportion of parental cells and not the final one. This customary choice is further reinforced by the fact that we have a less noisy estimate of initial proportions of cells than of the final, and so other definitions would lead to less precise measurements. Finally, our procedure can be viewed as standard competitive fitness assays but with initial ratio of the two types as a varied experimental parameter. Thus, for consistency with both theoretical and experimental literature, we associated the growth rates with the initial – more controlled – seeding proportion.

### F Generalizing the game assay to non-linear fitness functions

The game assay that we presented above is more interpretable and its output more easily plotted for matrix games with linear fitness functions. And in the case of our experimental system, the linear fitness functions provide an adequate fit for our purposes. However, that does not mean that the game assay has to be used only for linear games. Whereas section C used linear functions as the hypothesis class for fitting the growth-rate vs. proportion, one could use any other class of functions. An obvious candidate is polynomial fitness functions of orders higher than 1. We provide an example in Supplementary Figure 9 of a 3rd-order (cubic) fit. Visually, the cubic provides a better fit than the linear one in Figure 3, which is to be expected from the extra degrees of freedom. But qualitatively it provides the same interpretation as the linear fitness functions, including the same number of fixed points. In particular, DMSO + CAF has a single fixed point at *p* = 0.52 and (using the fixed region height from section C) a single fixed region for *p* ∈ (0.04, 0.668). This is much like the linear fit, but the fixed point region is expanded. The other three conditions (DMSO, Alectinib, and Alectinib + CAF) have no fixed points and no fixed regions.

**Supplementary Figure 10:**
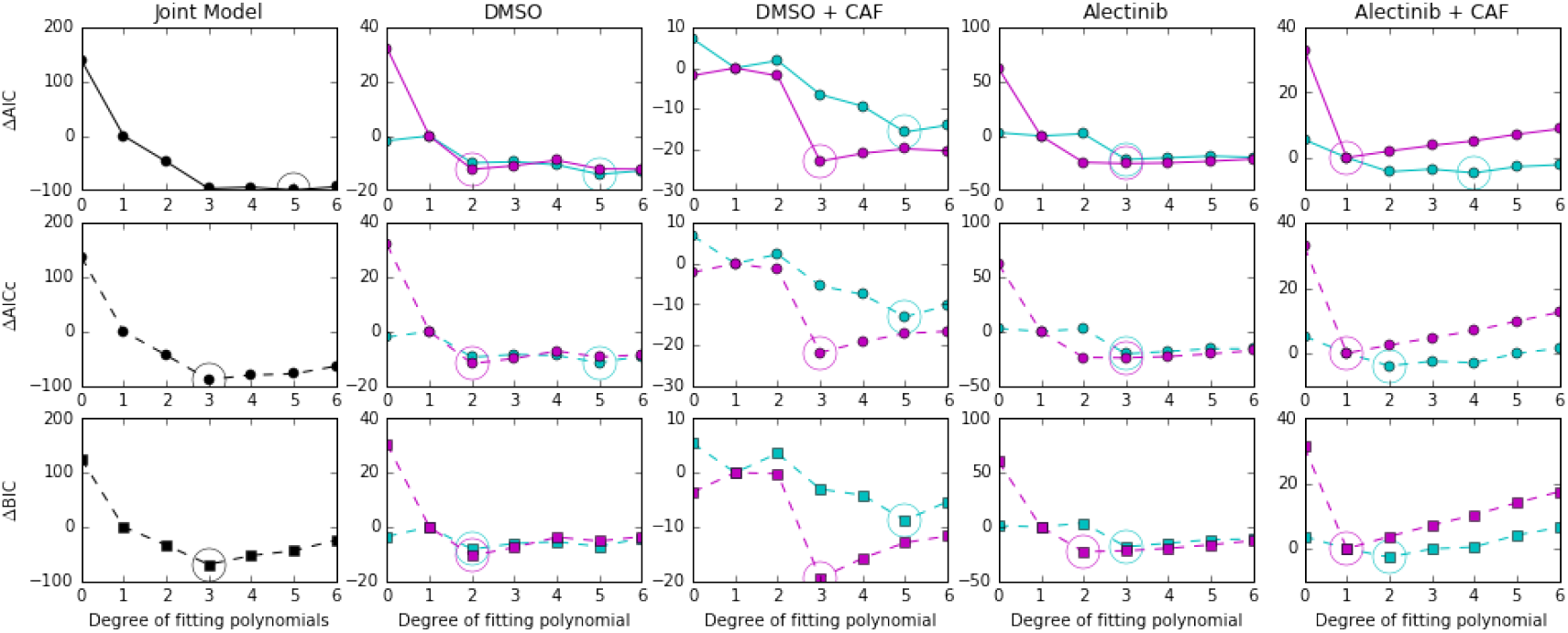
AICs and BICs for best polynomial fits of a given degree (up to additive offset). In cyan are AIC and BIC values for models parental cell fitness functions, while in magenta are AIC and BIC values for models of resistant cell fitness functions. Circles surround the minimum of AIC and BIC.

#### F.1 Information criteria for non-linear fits

If we treat the game assay not as a measurement and definition of games but as a model selection problem for parameter fitting then it becomes important to quantify the trade-off between the goodness of fit and model simplicity. For this, we can use techniques like the Akaike information criterion (AIC), its small-sample size correction (AICc), or the Bayesian information criterion (BIC) – or any other statistical model selection procedure. Given that (i) a polynomial of degree *d* has *k = d +* 2 degrees of freedom as a statistical model (+1 for zeroth order term, +1 for noise term); the eight models (4 conditions, 2 fitness functions per condition) are trained on *n* = 42 data points each; and AIC/BIC only works reasonablely when *n* >> *d*. For example, given our relatively small dataset for each model, Burnham & Anderson [46] would advocate to always prefer AICc over AIC (they suggest *n/k <* 40 as the cut off). Hence, we show the results of all three of AIC, AICc and BIC for polynomial fitness functions for degree *d ≤* 6 in Supplemental Figure 10. In this figure, a better model corresponds to a lower AIC, AICc or BIC value (lower on the y-axis). Since constant offsets in the information criteria do not matter for model selection, the axes are set so that the linear model has ∆{AIC, AICc, BIC} = 0. The leftmost column of Supplemental Figure 10 considers the joint product model where each fitness function has the same degree – for the *d* = 1 model, this would correspond to the linear game assay as presented in section C. Both AICc and BIC select the 3rd-degree polynomial model that we discussed above. AIC doesn’t differentiate strongly between the 3rd, 4th, 5th and 6th degree, but prefers slightly the 5th degree. Too much emphasis should not be placed on AIC however, given the number of parameters compared to sample size [46]. The four right columns of Supplemental Figure 10 consider independent models for each of the fitness functions across the 4 different conditions – so a total of 8 models. At the cost of extra researcher degrees of freedom, it is possible to look at the fits where the model for each of the 8 fitness functions is selected independently. Such a fit, as selected by BIC, is shown in Supplemental Figure 11. Note the two extra crossings of zero by the gain function in the DMSO + CAF case.

**Supplementary Figure 11:**
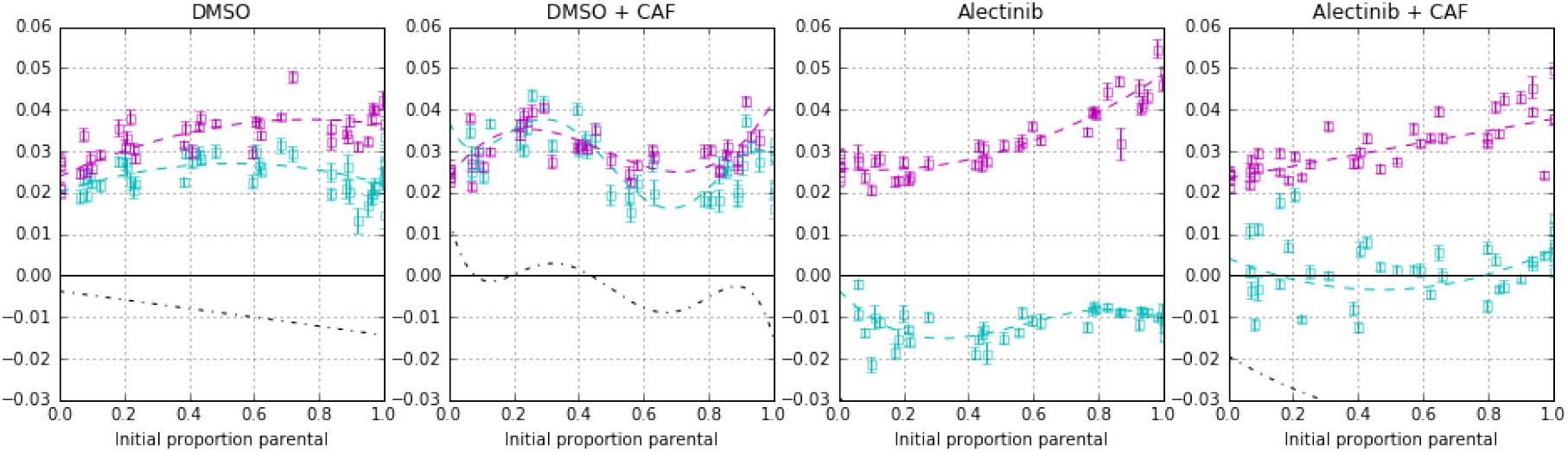
Fitness functions for competition of parental vs. resistant NSCLC as selected by BIC. For each plot: growth rate with confidence intervals versus initial proportion of parental cells. This is the same data as Figure 3. Cyan data points are growth rates of parental cells, and magenta for resistant cells. Dotted lines represent the fitness function of the least-squares best fit for models selected by BIC. These are a linear model for resistant fitness function in Alectinib + CAF; quadratic models for parental fitness function in Alectinib + CAF, resistant fitness function in Alectinib, and both fitness functions in DMSO; cubic for resistant in DMSO + CAF, and parental in Alectinib; and quintic for parental in DMSO + CAF. The black dotted line is the gain function for parental (see Figure 4a), it is identical with the *y* = 0 line in the DMSO + CAF condition (indication equal fitness for the two strategies) and well below it in the Alectinib conditions (indicating the strong advantage of resistance) and thus not visible in the figure.

**Supplementary Figure 12:**
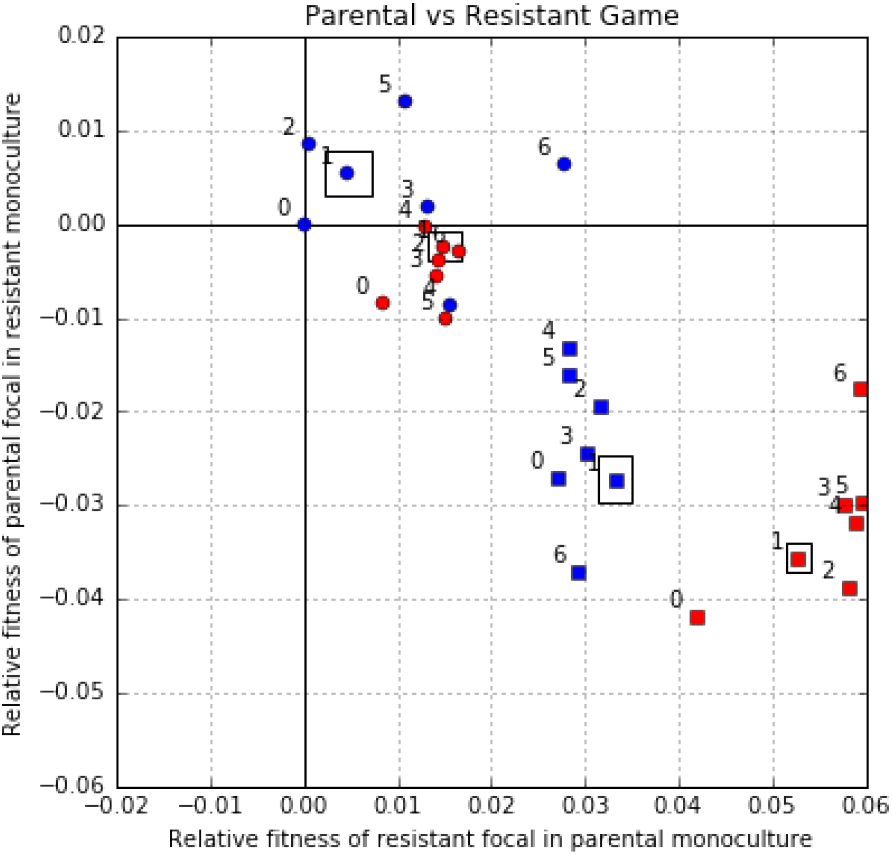
Mapping of the AIC and BIC selected fitness functions for the four conditions into game space. The x-axis is the relative fitness of a resistant focal in a parental monotypic culture. The y-axis is the relative fitness of a parental focal in a resistant monotypic culture. Games measured in our experimental system are specified by the bounding boxes corresponding to the range of their errors. The games corresponding to joint degree models are given as points, with joint degree labeled nearby. Experimental condition is represented by shape (DMSO: circle; Alectinib: square) and colour (no CAF: red; + CAF: blue).

#### F.2 Plotting nonlinear games

Just like with the linear games, it is possible to plot nonlinear games in our 2D game space based on the *p* = 0 and *p* = 1 endpoints of the gain function. We do this in Supplemental Figure 12 with each point labeled by the degree of the corresponding polynomial fitness functions. Unsurprisingly, at a brief glance there is broad qualitative agreement – all (but one) points are in the same quadrant as the linear model – although little quantitative agreement with the linear game assay – most points are outside of the error-box corresponding to the linear game. However for a general nonlinear game, unlike with linear games, two points in the same quadrant might not correspond to the same qualitative kind of dynamic. In particular, for a general nonlinear game, a quadrant only tells us the parity of the number of roots in (0,1) – where roots are counted by their multiplicity – and the order of alternations on the flow. For more on discrete flow alternation representation of gain functions, see Pena *et al*. [31].

Fortunately, for our particular experimental system the above generality is not realized. In particular, for all degrees of the DMSO, Alectinib, and Alectinib + CAF games the gain functions have no fixed points –just like the linear case. For DMSO + CAF, degrees *d* = {1,2,3} have one fixed point and *d* = {5, 6} have 3 fixed points (although only two fixed regions: *p* ∈ {(0.07, 0.25), (0.38, 0.46)} for *d* = 5 and *p* ∈ {(0.09, 0.31), (0.44, 0.50)} for *d* = 6. For *d* = 0 it is impossible for any model to be in the top right or bottom left quadrant – since no constant line can be both negative and positive – and there is no fixed point, but the fitness difference for DMSO is so tiny that there is a single fixed region for the whole space *p* ∈ (0,1). The real outlier for DMSO is *d* = 4 since it has two fixed points (and is thus in the bottom right quadrant) and two fixed regions at *p* ∈ {(0.07,0.11), (0.29,040)}. Thus, the existence of fixed point(s) in DMSO + CAF and absence of fixed points in the other conditions is robust across the nonlinear models. The exact position of the fixed point(s) in DMSO + CAF, however, is not as robust to model choice.

